# Patterns of gene expression and allele-specific expression vary among macular tissues and clinical stages of Age-related Macular Degeneration

**DOI:** 10.1101/2022.12.19.521092

**Authors:** Charles Zhang, Julie L. Barr, Leah A. Owen, Akbar Shakoor, Albert T. Vitale, John H Lillvis, Parker Cromwell, Nadine Husami, Robert Finley, Davis Ammar, Elizabeth Au, Neena B. Haider, Rylee A. Zavala, Elijah C. Graves, Mingyao Li, Amany Tawfik, Sarah X. Zhang, Dwight Stambolian, Michael H. Farkas, Ivana K. Kim, Richard M. Sherva, Lindsay A. Farrer, Margaret M. DeAngelis

## Abstract

Age-related macular degeneration (AMD) is a complex neurodegenerative disease and is the leading cause of blindness in the aging population. Early AMD is characterized by drusen in the macula and causes minimal changes in visual function. The later stages are responsible for the majority of visual impairment and blindness and can be either manifest as geographic atrophy (dry) or neovascular disease (wet). Available medicines are directed against the wet form and do not cure vision loss. Therefore, it is imperative to identify preventive and therapeutic targets. As the mechanism for AMD is unclear, we aim to interrogate the disease-affected tissue - the macular neural retina and macular retina pigment epithelium (RPE)/choroid. We investigated differentially expressed genes expression (DEG) across the clinical stages of AMD in meticulously dissected and phenotyped eyes using a standardized published protocol (Owen et al., 2019). Donor eyes (n=27) were obtained from Caucasian individuals with an age range of 60-94 and 63% were male, and tissue from the macula RPE/choroid and macula neural retina were taken from the same eye. Donor eyes were recovered within 6 hours post mortem interval time to ensure maximal preservation of RNA quality and accuracy of diagnosis. Eyes were then phenotyped by retina experts using multi modal imaging (fundus photos and SD-OCT). Utilizing DESeq2, followed PCA, Benjamini Hochberg adjustment to control for the false discovery rate, and Bonferonni correction for the number of paired comparisons: a total of 26,650 genes were expressed in the macula RPE/choroid and/or macula retina among which significant differential expression was found for 1,204 genes between neovascular AMD and normal eyes, 40 genes between intermediate AMD and normal eyes, and 1,194 genes between intermediate AMD and neovascular AMD. A comparison of intermediate AMD versus normal eyes included *TCN2, PON1, IFI6,* GPR123, and *TIMD4* as being some of the most significant DEGs in the macula RPE/choroid. A comparison of neovascular AMD versus normal eyes included *SLC1A2, SLC24A1, SCAMP5, PTPRN, and SEMA7A as* being some of the most significant DEGs in the macula RPE/choroid. Top pathways of DEGs in the macular RPE/choroid identified through Ingenuity Pathway Analysis (IPA) for the comparison of intermediate AMD with normal eyes were interferon signaling and Th1 and Th2 activation, while those for the comparison of neovascular AMD with normal eyes were the phototransduction and SNARE signaling pathways. Allele-specific expression (ASE) in coding regions of previously reported AMD risk loci identified by GWAS (Fritsche et al, 2016) revealed significant ASEs for C3 rs2230199 and CFH rs1061170 in the macula RPE/choroid for normal eyes and intermediate AMD, and for CFH rs1061147 in the macula RPE/choroid for normal eyes and intermediate and neovascular AMD. An investigation of the 34 established AMD risk loci revealed that 75% of them were significantly differentially expressed between normal macular RPE/choroid and macular neural retina, with 75% of these loci showing higher expression in the RPE. Similarly, disease state differences for the GWAS loci were only found to be statistically differentially expressed in the macular RPE/choroid. Moreover, the known coding variants in the previously identified GWAS loci including, *CFH*, *C3*, *CFB*, demonstrated ASE across AMD clinical stages in the macular RPE/choroid and not in the neural retina. These data at the bulk level underscore the importance of the RPE/choroid to AMD pathophysiology. While many bulk RNASeq data sets are publicly available, to the best of our knowledge this is one of the first publicly available datasets with both maculae RPE/choroid and macula neural retina from the same well phenotyped donor eye(s) where the macula is separated from the periphery. Our findings also underscore the importance of studying both macular tissue types to gain a full understanding of mechanisms leading to AMD. Our results provide insights into underlying biological mechanisms that may differentiate the disease subtypes and into the tissues affected by the disease.

## Introduction

Age-related macular degeneration (AMD), the leading cause of blindness in the aging population, is a complex neurodegenerative disease with both intermediate and late forms. The intermediate form is a clinical biomarker that can increase the risk of either of the two advanced forms; neovascular AMD (also referred to as wet AMD) and geographic atrophy (GA). In either form, this condition involves progressive degradation of the macula leading to central vision loss which impairs reading, facial recognition, and driving abilities [1]. Precise mechanisms contributing to disease pathogenesis and why the presence of disease is within the macula as opposed to peripheral regions of the retina remain unclear. Though AMD is phenotypically heterogeneous, the Age-Related Eye Disease Study (AREDS) classifies AMD into four stages based on the number and size of drusen in addition to observable pathological abnormalities, such as the sharp loss of pigmentation characteristic of geographic atrophy. These stages are denoted AREDS 1 (no AMD/normal aging), AREDS 2 (early AMD), AREDS 3 (intermediate AMD), and AREDS 4 (end-stage/advanced AMD) [2]. Tools used to assist the ophthalmologist in the staging of AMD include fundus imaging and spectral domain optical coherence tomography (SD-OCT) to provide direct visualization and/or high-resolution, non-invasive cross-sectional, enface imaging of the retina/RPE/choroid.

There is no cure for AMD. Neovascular AMD represents a smaller proportion of overall end stage AMD. The development of anti-vascular endothelial growth factor (VEGF) treatment has helped to mitigate visual loss associated with neovascular AMD, though cannot fully restore anatomic or visual integrity [3, 4]. There are no current FDA- approved therapeutic interventions for dry AMD which represents the majority of the AMD population. Currently the only treatment for intermediate AMD is supplementation of antioxidant AREDS2 formula, which has been demonstrated to modestly reduce the rate of progression to advanced AMD [6]. As yet there are no FDA approved medications for end stage dry AMD, geographic atrophy, although an anti-C3 and anti-C5 agent slowed the growth of geographic atrophy, both increased incidence of neovascular AMD; it is unclear when and if these agonists will be FDA approved [7, 8]. Appropriate therapies that prevent, slow or stop disease progression are clearly needed. Although a study by the International Age-related Macular Degeneration Genomics Consortium (IAMDGC) identified genome-wide significant association at 34 loci [3], most of the significantly associated variants are within non-coding regions (such as introns, gene regulatory regions, or positioned distally from a functional gene) and have no known functional consequences. While this has directed us to potential pathways underlying disease mechanism(s) this has not moved us closer to druggable therapeutic targets for AMD. Therefore, as we and others have previously hypothesized an investigation of well characterized and geographically dissected tissues affected by the disease pathophysiology may be the appropriate first step in ascribing function to a gene in a complex disorder such as AMD [9, 10]. Though gene expression has been examined within tissues affected by AMD, findings have not been consistent between studies [11–17]. Inconsistency between studies whether single cell, single nuclei and/or bulk RNASeq, may include lack of separation of geographical regions (eg. macula retina and/or the macula RPE/choroid from the periphery), a lengthy death to preservation time in the processing of the tissue, and/or lack of pre and/or post-mortem phenotyping for both eyes within the same donor [10,18, 20]. RNA-Seq is an agnostic and unbiased approach to examine gene expression a priori [21–22]. Additionally, RNA-Seq can uncover differentially spliced genes and long intervening non-coding RNAs (lncRNAs) [23–27]. To date, studies that have employed RNA-Seq [28–30] have not evaluated gene expression in both macula neural retina and macula RPE/choroid within the same donor eye in intermediate and separately neovascular AMD compared to well characterized control donor eyes. We focused the present study on tissues specifically affected by AMD, the macula of the retina pigment epithelium/choroid (RPE)/choroid and the macula of the neural retina, in an effort to ascertain differential gene expression (DEG) between the clinical stages of AMD. To address the complexity of a multi-faceted disease, we utilized a systems biology approach that integrates basic experimental, genomic, phenotypic, and clinical data into a model for understanding AMD disease pathogenesis. [28] We employed a standardized phenotyping protocol within a given donor, of both eyes recovered within 6 hours post mortem [20], the maximum interval to achieve minimal RNA half-life variability [29], enhanced RNA quality [34–36] and because we cannot assume that both normal and diseased tissue degrades at a similar rate. In addition, motivated by previous studies showing evidence of allele-specific expression (ASE) [37] in genes associated with risk of autism, stroke progression, Alzheimer disease and cancer [38–43], we interrogated the DNA of each donor for previously reported AMD GWAS coding variants [5] to determine whether an imbalance of expression between alleles may underlie phenotypic variation and hence the pathophysiology of AMD. To our knowledge, this is the first study to assess ASE across the clinical spectrum of AMD at a genome-wide level.

## Methods

The protocol was reviewed and approved by the institutional review board at the University at Utah (IRB 52879) and conforms to the tenets of the Declaration of Helsinki.

### Donor Eye Tissue Repository

Methods for human donor eye collection were previously described in detail according to a standardized protocol [20]. In brief, in collaboration with the Utah Lions Eye Bank, donor eyes were procured within a 6-hour post-mortem interval, defined as death-to-preservation time. Both eyes of the donor underwent post-mortem phenotyping with ocular imaging, including spectral domain optical coherence tomography (SD-OCT), and color fundus photography as published. Retinal pigment epithelium/choroid was immediately dissected from the overlying retina, and macula separated from periphery using an 8mm macular punch. For both peripheral and macular tissues, RPE/choroid was separated from the overlying retinal tissue using microdissection; tissue planes were optimized to minimize retinal contamination of RPE/choroid samples using a subsequent 6mm RPE/choroid tissue punch. Post-mortom phenotype was determined as published. In brief, AMD phenotype employed the <u>A</u>ge-<u>R</u>elated <u>E</u>ye <u>D</u>isease <u>S</u>tudy severity grading scale, where AREDS category 0/1 was considered normal, AREDS category 3 intermediate AMD, and AREDS category 4b neovascular AMD [44]. Phenotype analysis was performed as described, [24], by a team of 4 retinal specialists and ophthalmologists at the University of Utah School of Medicine, Moran Eye Center and the Massachusetts Eye and Ear Infirmary Retina Service. Agreement of all 4 specialists upon independent review of the color fundus and OCT imaging was deemed diagnostic; discrepancies were resolved by collaboration between a minimum of three specialists to ensure a robust and rigorous phenotypic analysis. One eye was per donor was biochemically analyzed. In the case of discordant phenotypes within the same donor, the more severe diseased eye was used for inclusion in the study. For example, if a patient had a diagnosis of AREDS 3 in one eye and AREDS 0/1 in the contralateral eye only the AREDS 3 eye was used in the study. Similarly, if a patient had a diagnosis of AREDS 3 in one eye and neovascular AMD in the contralateral eye only the neovascular eye was used in the study. Although AREDS category 2 early AMD, category 4a (geographic atrophy) and AREDS category 4c (both geographic atrophy and neovascular AMD) were collected and ascertained as previously described they were not included in this study [9].

### Study Population

Transcriptome profiling from macular retina and RPE/choroid samples from 27 unrelated eye tissue donors was performed using RNA-sequencing. Analyzed donor tissues included one randomly selected eye per donor comprising 10 eyes with intermediate AMD, 5 eyes with neovascular AMD, and 12 normal control eyes. Macular neural retina or RPE/choroid was isolated using the Utah Protocol technique as published by our group [20,45,-46]. In brief, neurosensory retina was isolated using an 8mm punch centered on the fovea and placed in RNAlater (Ambion). RPE/choroid tissue was isolated from overlying neurosensory retina using microdissection and retinal contamination minimized through isolation of the central 6mm; the tissue was similarly placed in RNAlater [20]. Tissues were stored at 4°C for 24 hours and then placed at −80° C for long term storage.

### Nucleic Acid Extraction and RNA-Sequencing

DNA and RNA were extracted from macular retina or RPE/choroid tissues prepared as above using the Qiagen All-prep DNA/RNA mini kit (cat #80204) according to manufacturer’s protocol (a total of 54 samples). Quality of RNA samples was assessed with an Agilent Bioanalyzer. Total RNA samples were poly-A selected and cDNA libraries were constructed using the Illumina TruSeq Stranded mRNA Sample Preparation Kit (cat# RS-122-2101, RS-122-2102) according to the manufacturer’s protocol. Sequencing libraries (18 pM) were chemically denatured and applied to an Illumina TruSeq v3 single read flow cell using an Illumina cBot. Hybridized molecules were clonally amplified and annealed to sequencing primers with reagents from an Illumina pTruSeq SR Cluster Kit v3-cBot-HS (GD-401-3001). Following transfer of the flowcell to an Illumina HiSeq instrument (HCS v2.0.12 and RTA v1.17.21.3), a multiplexed, 50 cycle single read sequence run was performed using TruSeq SBS v3 sequencing reagents (FC-401-3002).

### Primary Processing of RNA Sequencing Data

Each of the 54 sample 50bp, poly-A selected, non-stranded, Illumina HiSeq fastq datasets were processed as follows: reads were aligned using NovoCraft’s novoalign 2.08.03 software (http://www.novocraft.com/) with default settings plus the -o SAM -r All 50 options to output multiple repeat matches. The genome index used contained human hg19 chromosomes, phiX (an internal control), and all known and theoretical splice junctions based on Ensembl transcript annotations. Additional details for this aspect of the protocol are described elsewhere (http://useq.sourceforge.net/usageRNA-Seq.html). Next, raw novoalignments were processed using the open source USeq SamTranscriptiomeParser (http://useq.sourceforge.net) to remove alignments with an alignment score greater than 90 (∼ 3 mismatches), convert splice junction coordinates to genomic, and randomly select one alignment to represent reads that map equally well to multiple locations. Relative read coverage tracks were generated using the USeq Sam2USeq utility (http://useq.sourceforge.net/cmdLnMenus.html#Sam2USeq) for each sample and sample type (Normal Retina, Intermediate AMD Retina, Neovascular AMD Retina, Normal RPE/choroid, Intermediate AMD RPE, and Neovascular AMD RPE/choroid). These data tracks are directly comparable in genome browsers and good tools to visualize differential expression and splicing. Estimates of sample quality were determined by running the Picard CollectRNA-SeqMetrics application (http://broadinstitute.github.io/picard/) on each sample. These QC metrics were then merged into one spreadsheet to identify potential outliers. Agilent Bioanalyzer RNA integrity number (RIN) and library input concentration columns were similarly added for QC purposes (http://www.genomics.agilent.com). Out of the 54 samples, 48 samples passed QC analysis.

### Differential Gene Expression and Splicing Analysis

Sample sets were analyzed using the DefinedRegionDifferentialSeq (DRDS) utility of USeq to detect differentially expressed and differentially spliced genes. This application accepts as input a conditions directory containing folders with biological replicas from each sample type (Normal Retina, Intermediate AMD Retina, Neovascular AMD Retina, Normal RPE, Intermediate AMD RPE, and Neovascular AMD RPE) and an Ensembl gene table in UCSC refFlat format. Gene models were created by merging gene transcripts into a single composite “gene” with the USeq MergeUCSCGeneTable utility. A table containing alignment counts from each sample for each gene was created with DRDS. Data in this table provided the basis for estimating count-based differential abundance using the DESeq2 Bioconductor package (http://www.bioconductor.org/packages/release/bioc/html/DESeq2.html) [47]. This program estimates the over-dispersion in the count data and calculates p-values using a negative binomial test. The Benjamini-Hochberg p-value correction was applied to control for multiple testing. DESeq2 also generates a log2 ratio estimate of difference in gene abundance using variance corrected counts as well as rLog values for clustering and principal component analysis (PCA). Library size and within replica variance were estimated for each sample. Pairwise comparisons were made between the normal and disease subgroups. Differences in splicing were assessed for merged replica counts for each exon with ≥10 counts in each gene in each subgroup by a chi-square test. A Bonferroni multiple testing correction was applied and the exon with the biggest absolute log2 normalized gene count ratio was noted. A per base normalized gene count read coverage log2 ratio graph was created enabling visualization of the relative exon coverage difference for each pairwise comparison. To identify potential outlier samples, unsupervised hierarchical clustering (HC) and PCA were performed with the aid of the Partek Genomic Suite (http://www.partek.com/pgs) using the default settings. DESeq2 rLog values from genes with ≥20 counts were included in this procedure. For HC visualization, row values were mean centered at zero and scaled to a standard deviation of one.

### SNP Genotyping

Genome-wide SNP genotyping was performed on DNA obtained from peripheral blood from each donor eye subject using Illumina’s HumanOmni2.5-8 BeadChip Kit according to the manufacturer’s protocol (http://www.illumina.com/products/humanomni25-8_beadchip_kits.ilmn). AMD associated SNPs that were not present on the HumanOmni2.5-8 BeadChip were genotyped using a combination of pre-designed and Custom Taqman SNP Genotyping Assays (Applied Biosystems). Each assay was run in a 15 µl reaction containing 2x Taqman GTXpress master mix, 40x probe, and 10 ng of DNA. Thermal cycling was performed according to the manufacturer’s protocol. The ABI 7500 Real-Time PCR System, with the accompanying software, was used to analyze the genotypes.

### Differential Expression Validation by Real-Time PCR

RNA was reverse transcribed using oligo-dT primers (Invitrogen) and SuperScript III reverse transcriptase (Invitrogen) according to the manufacturer’s protocol. The cDNA was used as a template for real-time PCR reactions run in triplicate using pre-designed Taqman Gene Expression Assays (Life Technologies) for *UCHL1, PFKP, LPCAT1, PDPN, GAS1, CST3* and for *UBC* as an endogenous control. Assays were run on the Taqman 7500 Real Time PCR system (Life technologies). Mean Ct values were normalized to *UBC* and analyzed using REST 2009 Software (http://www.gene-quantification.de/rest.html).

### Allele-Specific Expression (ASE)

SNPs previously identified by GWAS as being associated with AMD (determined using the GWAS Catalog https://www.ebi.ac.uk/gwas/) [5] were investigated for allele-specific expression (ASE) in our dataset. Specifically, we genotyped the exonic AMD SNPs using either the genotypes from the HumanOmni2.5-8 BeadChip Kit or TaqMan assays. Bam files of individuals showing a heterozygous genotype were examined to determine the number of reads for each of the two alleles. Genotypes of heterozygotes determined from the SNP Chip showing monoallelic expression were confirmed using proxies (r^2^ ≥ 0.8) as determined by the 1000 Genomes phase 3 CEU reference panel. Only individuals with ≥ 10 reads were used. A binomial test, corrected using Benjamini-Hochberg, was used to determine statistically significant allelic imbalance within each individual.[48]

### Bioinformatic Analysis

QIAGEN Ingenuity Pathway Analysis (IPA) (QIAGEN Inc., https://digitalinsights.qiagen.com/IPA) was employed to identify pathways our DEGs function in between AMD normal, Intermediate AMD, and Neovascular AMD [49]. Identification of potential pathways associated with protective or risk features was conducted between the neovascular AMD vs intermediate AMD through direction of log fold changes showing higher expression in tissue associated with either intermediate AMD or neovascular AMD. Genes with higher expression with intermediate AMD were labelled as protective while higher expression in neovascular AMD. The list of protective and risk genes were compared against the genes associated with the significant canonical pathways identified through IPA.

## Results

Numbers and characteristics of subjects available for analysis of each tissue type after QC are shown in **Table 1**. For clustering analysis, using both hierarchal clustering (HCA) and principal component analysis (PCA) based on the samples’ whole transcriptome expression, samples split into two primary groups comprised of Retina and RPE/choroid samples. Clear separation of RPE/choroid and retina tissue types was observed. Within each tissue type, there was not clear separation between the disease subtypes. Greater variability was observed among the RPE/choroid samples than among the retina samples (**Fig 1**). None of the samples differed substantially to warrant flagging as an outlier.

**Fig 1.**
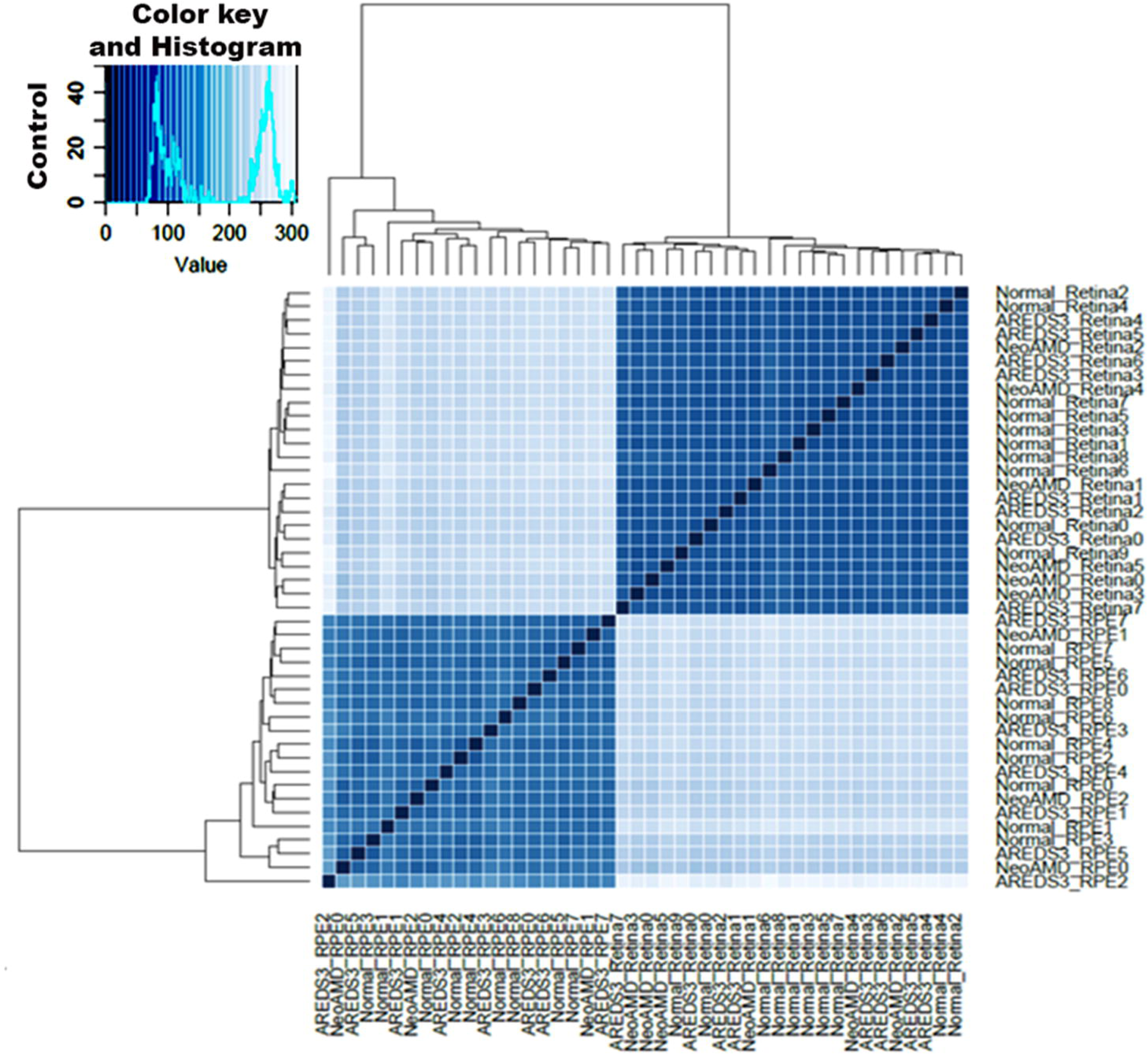
Hierarchal clustering analysis (HCA) of 47 donor eye samples from the RPE/choroid and retina. Rows and columns represent each individual sample against the rest of the samples.

**Table 1.** Subject Characteristics **Abbreviations:** n, number; Ave., average; RIN, RNA Integrity Number; RPE, Retinal Pigment Epithelium.

| Normal |  |  |  |  |
| --- | --- | --- | --- | --- |
| Group | n | Ave. RIN | Age (range) | Males (%) |
| All Samples (Retina) | 12 | 6.65 | 74.0 (60-94) | 9 (75.0%) |
| All Samples (RPE/Choroid) | 12 | 6.66 | 74.0 (60-94) | 9 (75.0%) |
| Clean Samples (Retina) | 10 | 6.76 | 74.4 (60-94) | 8 (80.0%) |
| Clean Samples (RPE/Choroid) | 9 | 6.93 | 74.2 (60-94) | 7 (77.8%) |
| Intermediate AMD |  |  |  |  |
| Group | n | Ave. RIN | Age (range) | Males (%) |
| All Samples (Retina) | 10 | 6.89 | 76.0 (60-87) | 6 (60.0%) |
| All Samples (RPE/Choroid) | 10 | 6.7 | 76.0 (60-87) | 6 (60.0%) |
| Clean Samples (Retina) | 9 | 6.91 | 75.0 (60-87) | 6 (66.7%) |
| Clean Samples (RPE/Choroid) | 9 | 6.76 | 75.0 (60-87) | 7 (66.7%) |
| Neovascular AMD |  |  |  |  |
| Group | n | Ave. RIN | Age (range) | Males (%) |
| All Samples (Retina) | 5 | 6.7 | 83.4 (74-94) | 2 (40.0%) |
| All Samples (RPE/Choroid) | 5 | 7.06 | 83.4 (74-94) | 2 (40.0%) |
| Clean Samples (Retina) | 5 | 6.7 | 83.4 (74-94) | 2 (40.0%) |
| Clean Samples (RPE/Choroid) | 5 | 7.06 | 83.4 (74-94) | 2 (40.0%) |

To evaluate the quality of our tissue dissection we calculated the number of reads mapped to genes known to be expressed exclusively in the neural retina and RPE/choroid, respectively, using an approach as previously described for the retina [61]. Retina genes involved in phototransduction (*GNGT1, GUCA1A, PDE6A, GNB1, CNGB1, GNAT1, CNGA1, PDE6B, PDE6G, PRPH2, RHO, ROM1, SAG*, and *SLC24A1*) accounted for an average of 2.3% of reads in the total normal retina library and accounted for 0.06% of our normal RPE/choroid tissue reads, proportions which are similar to those reported in a previous study [50]. In our study RPE/choroid genes (*BEST1, RDH5*, and *RPE6*,) accounted for an average of 0.65% of reads in the total RPE/choroid library and only 0.02% of total reads in the retina library. These findings demonstrate that neither the retina or the RPE/choroid is likely contaminated (e.g., if there was contamination of retina genes in the RPE/choroid library, reads would be greater than 1% compared to the observed proportion of 0.06%).

### Gene Expression Differences

A total of 26,650 genes were expressed in RPE/choroid and/or retina. Within phenotypically documented normal eyes, 16,638 genes showed significant (FDR ≤ 0.05) differential expression between RPE/choroid and retina tissues with a minimum fold change ≥ |1.5|. Within macular RPE/choroid tissues, significant differential expression was observed for 1,204 genes between neovascular AMD and normal eyes, 40 genes between intermediate AMD (AREDS 3) and normal eyes, and 1,194 genes between Intermediate AMD and neovascular AMD eyes (**Fig 2a; S1 Table**). Within macular neural retina tissues, 41 genes were differentially expressed between neovascular AMD and normal eyes, 30 genes between Intermediate AMD and normal eyes, and 50 genes between Intermediate AMD and neovascular AMD eyes (**Fig 2b**). Of these differentially expressed genes, 29 were unique to Intermediate AMD vs. normal RPE/choroid, 285 were unique to neovascular AMD vs. normal, and 276 were unique to intermediate AMD vs. neovascular AMD. Of the 40 significant differentially expressed genes in the normal versus Intermediate AMD RPE/choroid and the 1204 significant differentially expressed genes in the normal versus neovascular AMD RPE/choroid, six genes found in common (**Fig 3**). These six common genes were *MTRN2L1, CLEC2L, CCM2L, CYP4X1, GLDN,* and *SMAD7* (**Fig 3**). The top 10 differentially expressed genes between disease states within the RPE/choroid and Retina are listed in **Table 2a-b**. Of note, the only gene, that overlapped between any RPE/choroid and retina disease comparisons, was mitochondrial-derived peptide humanin (*MTRNR2L1*). *MTRNR2L1* was found to be differentially expressed between the intermediate AMD versus normal state. Of interest in the context of AMD, *MTRNR2L1* has been previously reported to protect human RPE cells against oxidative stress and restore mitochondrial function *in vitro* [51]. *CHD7* which was found to be downregulated in those with intermediate AMD compared to normal RPE/choroid has been previously associated with AMD in non-smokers by GWAS [52]. *TIMD4*, upregulated in those with intermediate AMD compared to normal RPE is associated with regulation of blood level lipids (*LDL*, cholesterol and triglycerides) at the GWAS level [53–55]. Also noteworthy, a unique lncRNA (*AC000124.1*) was down regulated in intermediate AMD compared to neovascular RPE/choroid, while lincRNA (*RP11-240M16.1*) and *PIWL1* were both upregulated in neovascular AMD compared to normal RPE/choroid (Tables 2 and 2b, respectively). A total of nine unique miRNAs were identified (*MIR146A*, *MIR3918*, *MIR4657*, *MIR17HG*, *MIR3620*, *MIR 3064*, *MIR197*, *MIR4680*, and *MIR4647*) across all disease comparisons. Of these miRNAs, six (*MIR4657*, *MIR17HG*, *MIR3620*, *MIR197*, *MIR3064*, and *MIR3918*) were found to have DEG in intermediate AMD vs neovascular AMD in the RPE/choroid while three (*MIR146A*, *MIR197*, and *MIR3918*) were identified in the neovascular AMD vs normal in the RPE/choroid (**Table S1**). In the retina, two miRNAs were seen, *MIR4680,* in the neovascular vs normal and *MIR4647* in the intermediate AMD vs neovascular AMD (**Table S1**).

**Fig 2.**
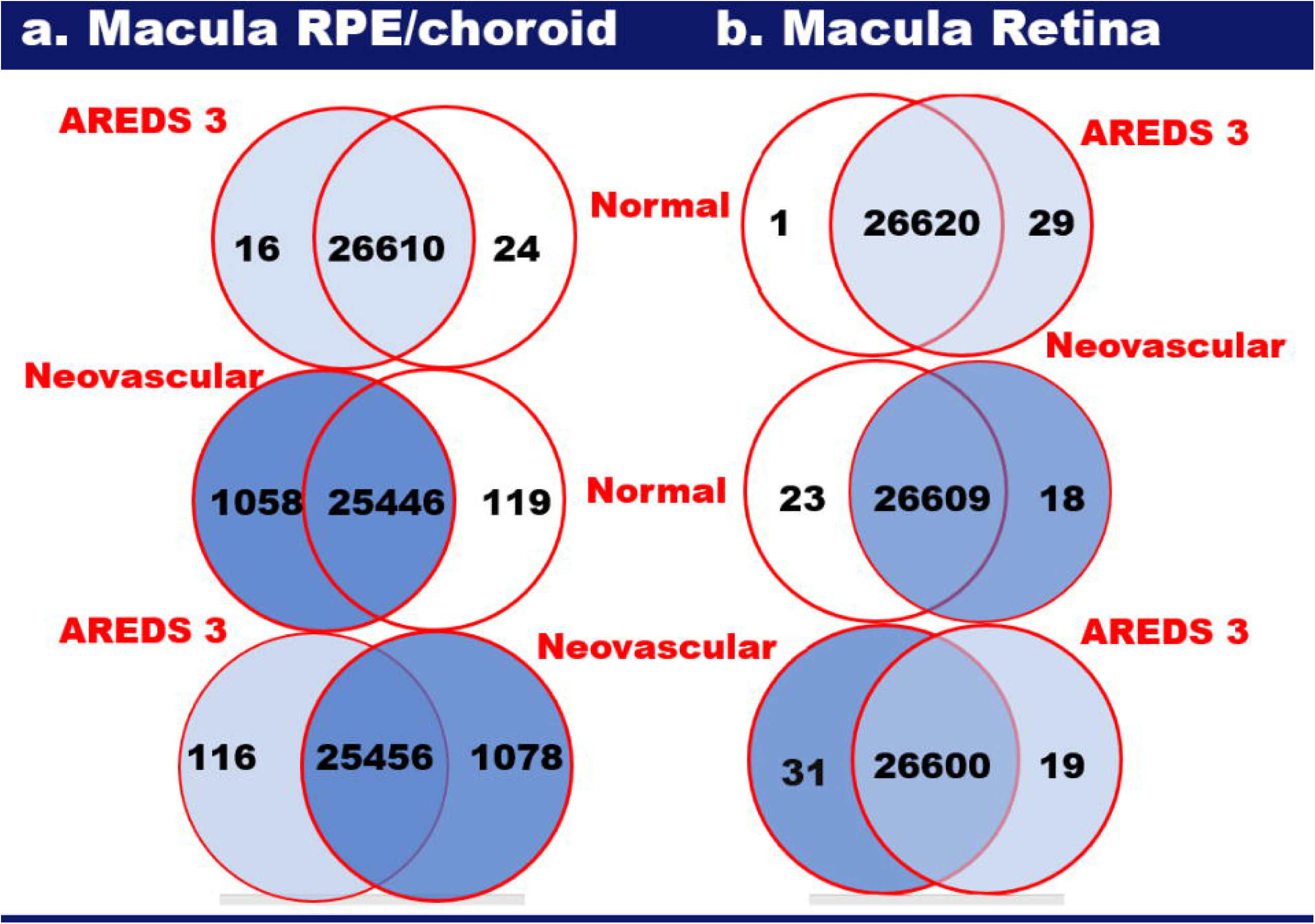
Differentially expressed genes. Each circle shows the number of genes with increased expression in that disease state, i.e., 1085 genes were upregulated in the macula RPE/choroid of neovascular eyes when compared to normal eyes and 119 genes were upregulated in the macula RPE/choroid of normal eyes compared to neovascular eyes.

**Fig 3.**
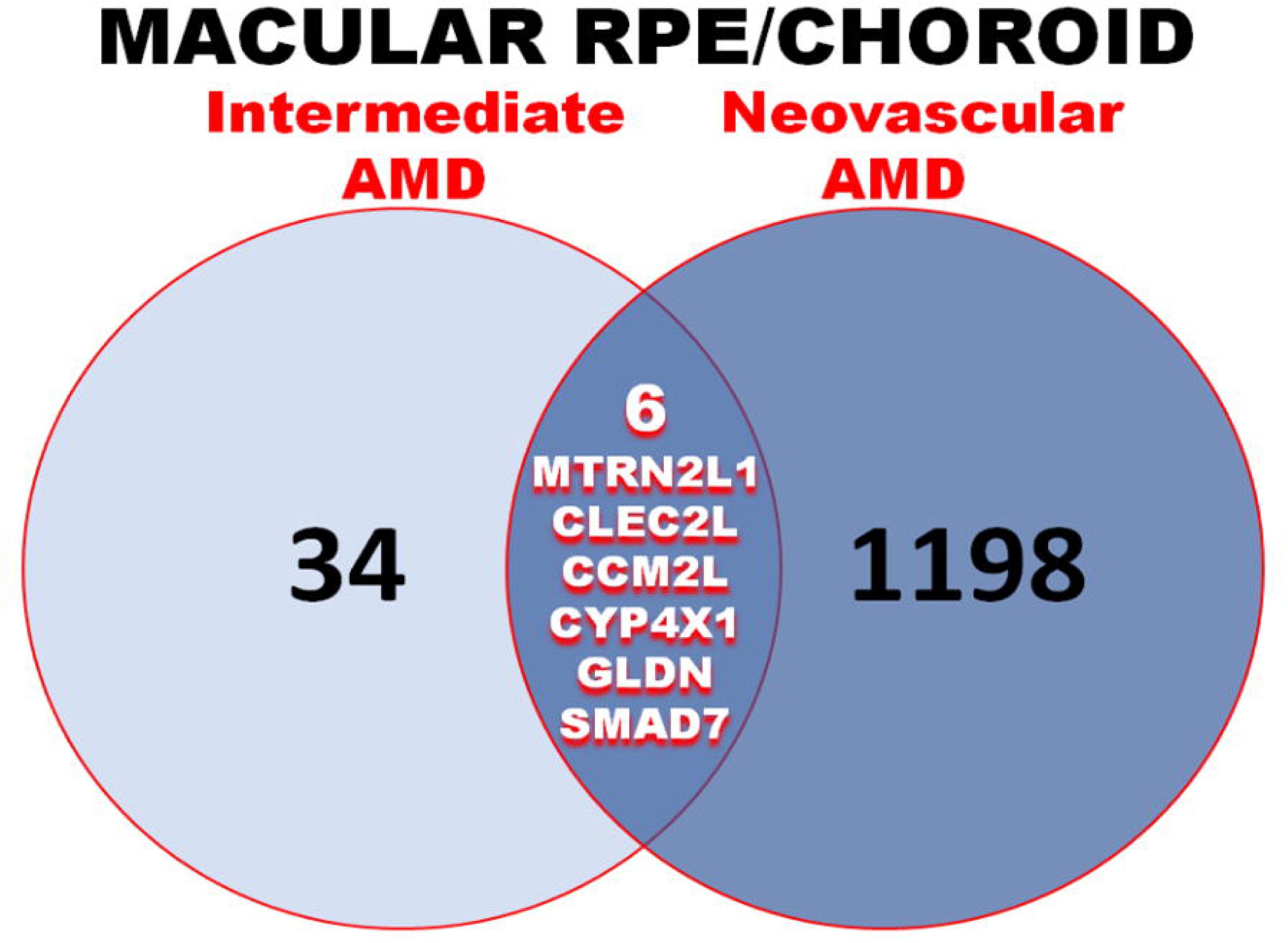
Overlap of differentially expressed genes. Each circle represents the number of significant differentially expressed genes in RPE/choroid when comparing normal eyes with both neovascular AMD and AREDS3. The overlap between these two circles shows the number of common genes between these two conditions.

**Table 2.**
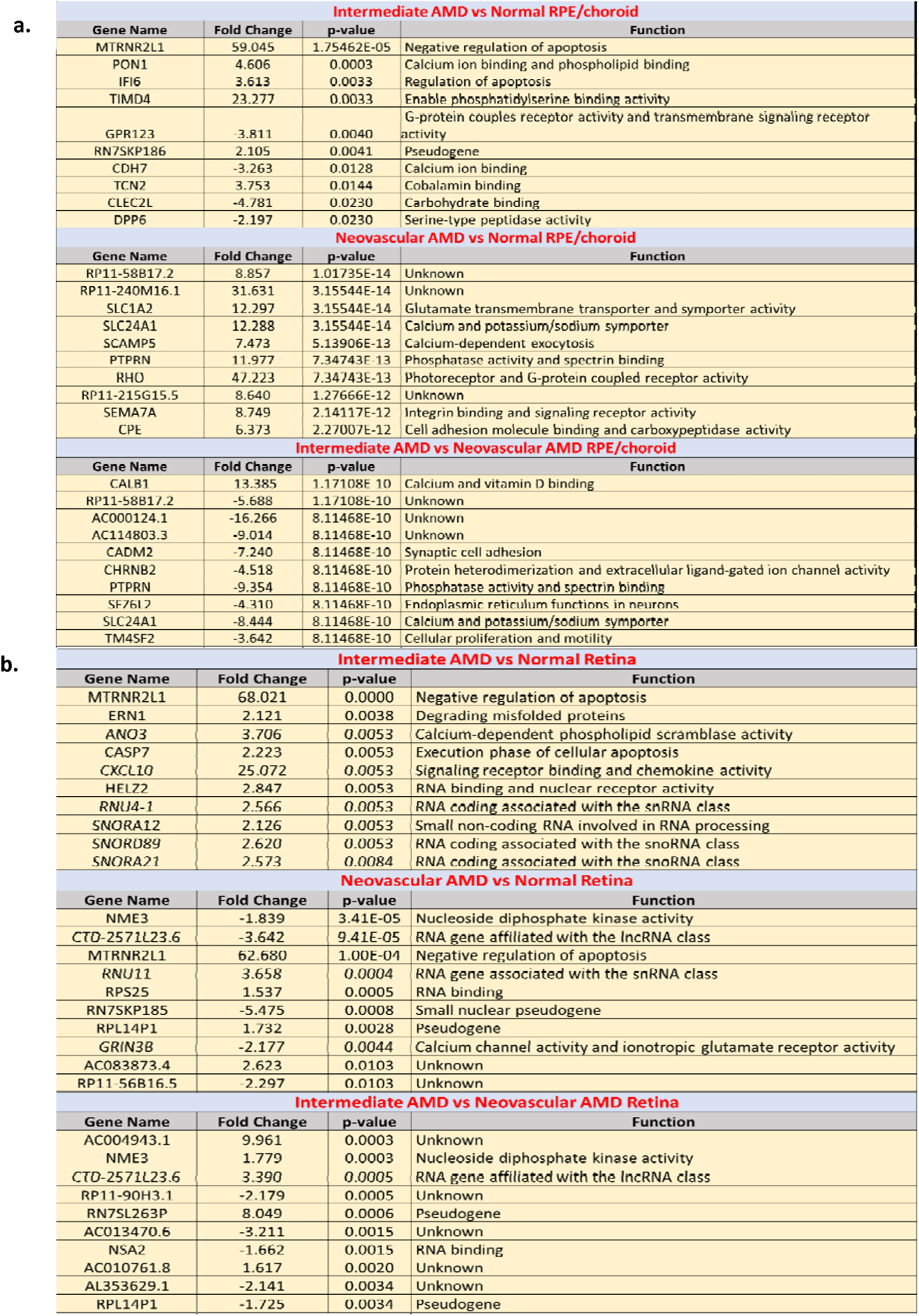
Top ten significant differentially expressed genes in each disease comparison.

### Analysis of Genes previously associated with AMD

Among genes demonstrated to be associated with AMD risk in previous studies [10,56–57] and the GWAS conducted by the International AMD Genetics Consortium (IAMDGC) [1], expression was statistically significantly *higher* for *ABCA4* (3.1 fold increase, p-value = 0.0001), *ABCA7* (3.4 fold increase, p-value = 0.0018), and *RORA* (1.9 fold increase, p-value = 0.004) in neovascular RPE/choroid tissue compared with normal RPE/choroid tissue (fold change > 2, p-value ≤ 0.01; **Table S1**). *VTN* expression (fold change = 3.3, p-value = 0.02) was also found to be significantly *higher* in neovascular RPE/choroid tissue compared with normal RPE/choroid tissue. Consistent with these results, we found that expression of *ABCA4* (fold change = −3.1, p-value = 0.0002) and *ABCA7* (fold change = −3.2, p-value = 0.0018) was significantly *lower* in intermediate AMD RPE/choroid tissue compared with neovascular RPE/choroid tissue. (fold change > 2, p-value ≤ 0.01). Expression was also found to be significantly *higher* for *TNFRSF10B* (fold change = 1.7, p-value = 0.02) and *TRPM1* (fold change = 1.7, p-value = 0.04) in intermediate AMD RPE/choroid tissue compared with neovascular RPE/choroid tissue. Additionally, *SPEF2* expression (fold change = −1.5, p-value = 0.0165) was significantly *lower* in intermediate AMD RPE/choroid tissues compared to neovascular RPE/choroid tissue. None of the associated AMD risk susceptibility genes were significantly differentially expressed between intermediate AMD and normal RPE/choroid tissues.

Within normal macular tissues, the expression levels of *ABCA4, ABCA7, C10orf88, CYP24A1, NLRP2, PELI3, RORA, RORB, SPEF2, SYN3, TMEM97,* and *VTN* were significantly higher in the neural retina compared to the RPE/choroid, whereas the levels of *ABCA1, ABHD2, ADAMTS9-AS1, ADAMTS9-AS2, C2, C3, C4A, C9*, *CD63, CETP, CFB, CFH, CFHR3-CFHR1, CFI, CNN2, COL5A1, COL8A1, ITGA7, LIPC, MMP19, MMP9, PCOLCE, PILRA, PKP2, PLA2G4A, RDH5, RGS13, RLBP1, SLC16A8, TGFBR1, TIMP3, TNFRSF10A, TNFRSF10B, TRPM1, TRPM3, TSPAN10, UNC93B1,* and *VDR* were significantly higher in the RPE/choroid compared to the retina (threshold: fold change > 2, p-value ≤ 0.01; **Table S1**). *ARHGAP21, CCT3, HTRA1,* and *SRPK2* were also found to be up-regulated in the neural retina compared to the RPE/choroid, whereas *NPLOC4, RAD51B, ROBO1, SKIV2L,* and *STON1-GTF2A1L* were found to be marginally up-regulated in the RPE/choroid compared to the neural retina (**Table S1**). Overall, of the 34 GWAS loci (5), 75% were found to be statistically differentially expressed between the macular Retina and the macular RPE/choroid, of these genes 75% were significantly upregulated in the RPE/choroid (**Table 3**).

**Table 3.** Comparisons of tissue expression levels in RPE/Choroid or retina of previously established AMD risk loci among normal RPE/choroid vs normal retina eyes.

| Previously Identified AMD Loci | Tissue With Higher Expression (RPE/Choroid or Retina) |
| --- | --- |
| C2/CFB/SKIV2L | RPE/Choroid |
| ARMS2/HTRA1 | Retina |
| SYN3 * | Retina |
| TIMP3 * | RPE/Choroid |
| PRLR/SPEF2 | RPE/Choroid |
| PILRB/PILRA | RPE/Choroid |
| KMT2E/SRPK2 | Retina |
| MIR6130/RORB | RPE/Choroid |
| RDH5/CD63 | RPE/Choroid |
| CTRB2/CTRB1 | RPE/Choroid |
| TMEM97/VTN | Retina |
| NPLOC/TSPAN10 | RPE/Choroid |
| ABCA1 | RPE/Choroid |
| ADAMTS9-AS2 | RPE/Choroid |
| COL8A1 | RPE/Choroid |
| C9 | RPE/Choroid |
| TNFRSF10A | RPE/Choroid |
| TGFRB1 | RPE/Choroid |
| RAD51B | RPE/Choroid |
| LIPC | RPE/Choroid |
| CETP | RPE/Choroid |
| C3 | RPE/Choroid |
| SLC16A8 | RPE/Choroid |
| TRPM3 | RPE/Choroid |
| RORB | Retina |
| ARHGAP21 | Retina |
| CNN2 | RPE/Choroid |
| MMP9 | RPE/Choroid |

Table 3. Shows tissue with higher expression of previously identified AMD loci as identified by Fritsche et al. (2016) for the RPE/Choroid versus the retina in normal eyes.

*Identified on the same locus but have differing tissue with higher expression

### Pathway Analysis for Differentially Expressed Genes

IPA pathway analysis was conducted on significant differentially expressed genes found in the RPE/choroid across all disease states. For each comparison, the ten most significant pathways for each condition were identified (**Table 4**). When comparing intermediate AMD and normal, 21 significant pathways for significant differentially expressed genes were revealed (**Table 4**). A total of 63 significant pathways were identified when looking into significant differentially expressed genes in neovascular AMD with normal, and 52 significant pathways when comparing neovascular AMD with intermediate AMD (**Table 4**). Interestingly, the calcium signaling pathway was significant across all comparisons, while 47 pathways were in common between the neovascular AMD and normal with the neovascular and intermediate AMD (**Table 4**). A pathway analysis of the six overlapping genes between intermediate AMD and normal with neovascular AMD and normal revealed 14 significant pathways, with BMP signaling and TGF-β signaling being of particular interest. The axonal guidance signaling pathway showed potential protective features with in the intermediate AMD vs neovascular AMD in the RPE/choroid 23% (*ACE2*, *ADAMTS1*, *ADAMTS8*, *BMP8A*, *RND1*, *SEMA4C*, and *WNT16*) of the identified genes being protective in this pathway while 10% (*EPHA10*, *LRRC4C*, and *SEMA4B*) shown as risk genes with the remaining 67% of genes in this pathway not being identified.

**Table 4.** IPA pathway analysis of significant differentially expressed genes in the RPE/Choroid.

| Intermediate AMD vs Normal |  | Neovascular AMD vs Normal |  | Neovascular AMD vs Intermediate AMD |  |
| --- | --- | --- | --- | --- | --- |
| Pathway Name | P-Value | Pathway Name | P-Value | Pathway Name | P-Value |
| Interferon Signaling | 2.68E-05 | Phototransduction Pathway | 1.08E-29 | Phototransduction Pathway | 8.41E-30 |
| Th1 and Th2 Activation Pathway | 1.68E-04 | SNARE Signaling Pathway | 3.37E-07 | SNARE Signaling Pathway | 6.06E-08 |
| Th1 Pathway | 1.01E-03 | Cardiac β-adrenergic Signaling | 1.72E-06 | Cardiac β-adrenergic Signaling | 1.50E-06 |
| nNOS Signaling in Neurons | 2.62E-03 | G-Protein Coupled Receptor Signaling | 5.51E-06 | Synaptogenesis Signaling Pathway | 2.45E-06 |
| Role of JAK2 in Hormone-like Cytokine Signaling | 4.52E-03 | Endocannabinoid Neuronal Synapse Pathway | 5.69E-06 | Protein Kinase A Signaling | 5.95E-05 |
| Activation of IRF by Cytosolic Pattern Recognition Receptors | 4.96E-03 | Role of NFAT in Cardiac Hypertrophy | 5.04E-05 | GABA Receptor Signaling | 6.26E-05 |
| Multiple Sclerosis Signaling Pathway | 4.58E-03 | Synaptogenesis Signaling Pathway | 6.10E-05 | Endocannabinoid Neuronal Synapse Pathway | 7.18E-05 |
| JAK/STAT Signaling | 7.78E-03 | GABA Receptor Signaling | 6.88E-05 | G-Protein Coupled Receptor Signaling | 3.81E-04 |
| Role of Hypercytokinemia/hyperchemokinaemia in the Pathogenesis of Influenza | 8.53E-03 | Protein Kinase A Signaling | 6.95E-05 | Role of NFAT in Cardiac Hypertrophy | 3.69E-04 |
| IL-15 Production | 1.69E-02 | Relaxin Signaling | 1.27E-04 | Relaxin Signaling | 3.76E-04 |

Table 4. Shows the top ten most significant pathways of the significant differentially expressed genes across all disease conditions. A comparison of the significant pathways in two comparisons, Neovascular AMD vs Normal and Neovascular AMD vs Intermediate AMD with the top ten shown for each comparison.

### Gene splicing differences among AMD stages and tissue types

Differential splicing was observed for 1,154 genes within the retina and for 629 genes within the RPE/choroid in a comparison of neovascular AMD and normal tissues (**Fig 4**). Fewer differentially spliced genes were observed for comparisons of intermediate AMD vs. neovascular AMD (810 in retina, 608 in RPE/choroid) and intermediate AMD vs. normal (210 in retina, 177 in RPE/choroid). Only a small proportion of the 629 differentially spliced genes in the RPE choroid were also differentially expressed in group comparisons: 113 neovascular AMD vs. normal (18.0%), 98 intermediate AMD vs. neovascular AMD (16.1%), and 6 intermediate AMD vs. normal (3.4%). There were no overlapping differentially expressed and spliced genes within the retina. A much larger number of genes in the retina were differentially spliced than in the RPE for all group comparisons.

**Fig 4.**
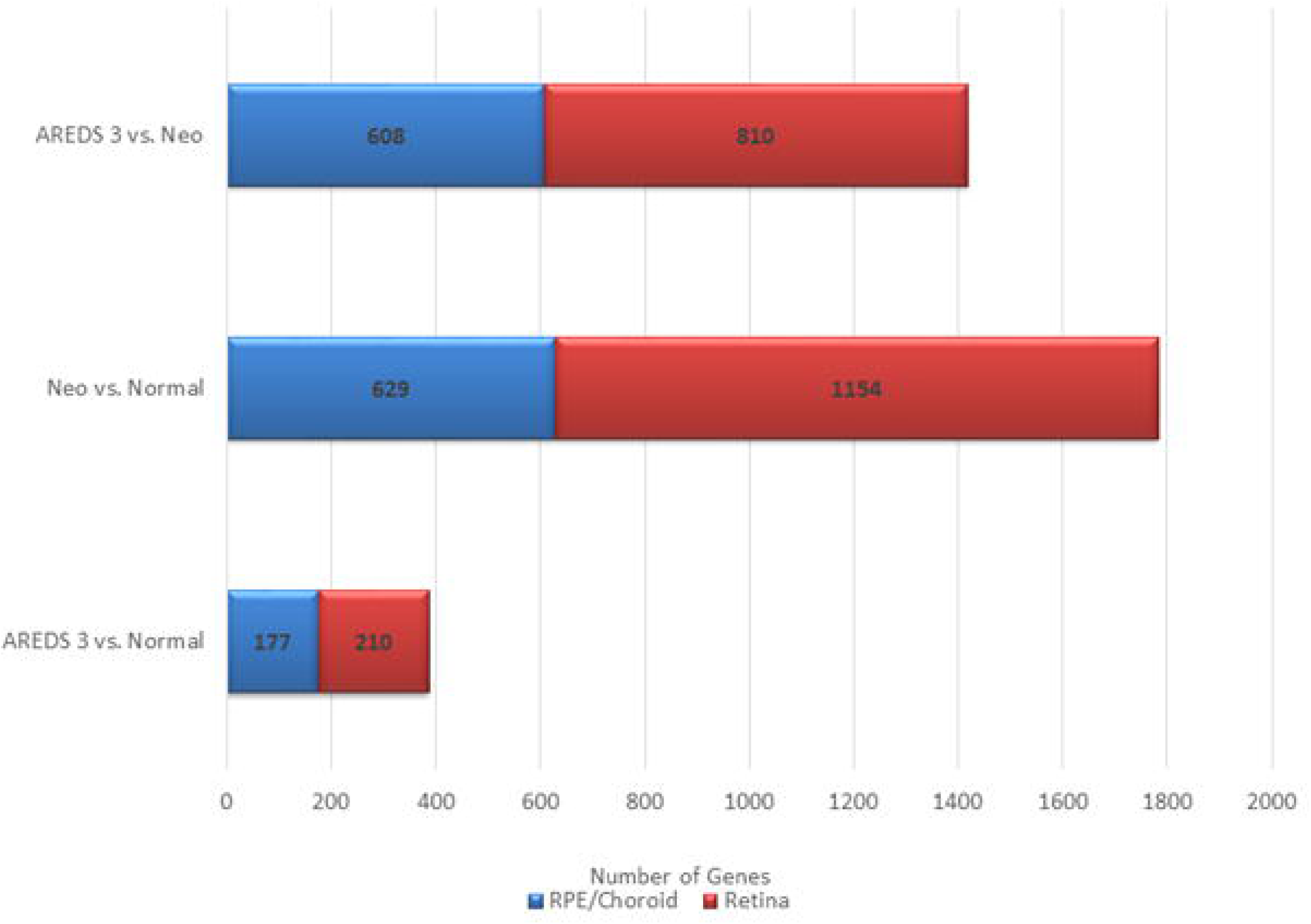
Differentially spliced genes. The number of genes showing differential splice sites among each of the comparisons examined.

### Allele Specific Expression (ASE)

According to annotation information for published AMD genome-wide association studies included in the NHGRI-EBI Catalog (https://www.ebi.ac.uk/gwas/home; accessed 3/28/17), 12 AMD-associated SNPs are located in coding regions of *APOE* (1), *ARMS2* (1), *C2* (1), *C3* (2), *CFB* (1), *CFH* (4), *CFI* (1), and *PLA2G12A* (1), and therefore were investigated for allele specific expression (**Table 5**). No heterozygotes were found in our sample for *CFH* rs121913059, *CFI* rs141853578, *C3* rs147859257, or *APOE* rs429358. We found no expression of *ARMS2* in either the RPE/choroid or neural retina and therefore we could not investigate the coding SNP rs10490924. For those heterozygotes showing mono-allelic expression (n=6), *CFH* rs10754199 was used to confirm heterozygotes for *CFH* coding SNPs rs10661170 and rs1061147 (r2 = 1 for rs10754199 and both coding SNPs), *CFB* rs2242572 was used to confirm heterozygous genotypes for *CFB* rs641153 (r2 = 1), and *C3* rs1047286 was used to confirm the heterozygous genotype of *C3* rs2230199 (r2 = 0.843). Significant ASE was detected within individuals at 4 SNPs: *CFH* rs1061170 (Y402H), *CFH* rs1061147, *CFB* rs641153, and *C3* rs2230199. Specifically, for *CFH* rs1061170 we found significant ASE within 2/6 intermediate AMD RPE samples, and 1/7 normal RPE samples. None of the four neovascular AMD RPE heterozygotes showed ASE, noting that there were 10 or fewer reads for these samples in the retina data. For rs1061147, significant ASE was observed within 5/6 intermediate AMD RPE samples, 3/4 neovascular RPE samples, and 7/7 normal RPE samples. These same heterozygotes had 10 or fewer reads among the retina data. The single heterozygote for *CFB* rs641153 (a normal sample) showed significant ASE within the RPE tissue. There were 10 or fewer reads for this SNP in macula retina. There was significant ASE for *C3* rs2230199 within 2/3 intermediate AMD RPE samples, 0/1 neovascular AMD RPE samples, and 2/2 normal RPE samples. These same heterozygotes had 10 or fewer reads in retina tissue.

**Table 5.** Allele Specific Expression (ASE) of known AMD associated SNPs. **Abbreviations:** *, Only individuals with greater than 10 reads are counted; SNP, Single Nucleotide Polymorphism; Hets, heterozygotes; RPE, Retinal Pigment Epithelium.

| SNP | Location | #Hets | Individuals with Significant ASE ($p < .05$)* | | | | | |
| --- | --- | --- | --- | --- | --- | --- | --- | --- |
|  |  |  | Intermediate AMD Retina | Neovascular Retina | Normal Retina | Intermediate AMD RPE | Neovascular RPE | Normal RPE |
| CFH rs1061147 | chr1:196654324 | 18 | 0/0 | 0/0 | 0/0 | 5/6 | 3/4 | 7/7 |
| CFH rs1061170 | chr1:196659237 | 18 | 0/0 | 0/0 | 0/0 | 2/6 | 0/4 | 1/7 |
| CFH rs35292876 | chr1:196706642 | 1 | 0/0 | 0/0 | 0/0 | 0/1 | 0/0 | 0/0 |
| CFH rs121913059 | chr1:196716375 | 0 | 0/0 | 0/0 | 0/0 | 0/0 | 0/0 | 0/0 |
| PLA2G4A rs2285714 | chr4:110638810 | 15 | 0/1 | 0/1 | 0/3 | 0/3 | 0/0 | 0/1 |
| CFI rs141853578 | chr4:110685820 | 0 | 0/0 | 0/0 | 0/0 | 0/0 | 0/0 | 0/0 |
| C2 rs9332739 | chr6:31903804 | 4 | 0/0 | 0/0 | 0/0 | 0/1 | 0/0 | 0/2 |
| CFB rs641153 | chr6:31914180 | 6 | 0/0 | 0/0 | 0/0 | 0/0 | 0/0 | 1/1 |
| ARMS2 rs10490924 | chr10:124214448 | 7 | 0/0 | 0/0 | 0/0 | 0/0 | 0/0 | 0/0 |
| APOE rs429358 | chr19:45411941 | 0 | 0/0 | 0/0 | 0/0 | 0/0 | 0/0 | 0/0 |
| C3 rs147859257 | chr19:6718146 | 0 | 0/0 | 0/0 | 0/0 | 0/0 | 0/0 | 0/0 |
| C3 rs2230199 | chr19:6718387 | 6 | 0/0 | 0/0 | 0/0 | 2/3 | 0/1 | 2/2 |

### Validation and replication of RNA-Seq findings

We validated our RNA-Seq methodology by choosing genes that varied in fold expression from a range of +20 to −20 (FDR of p < 0.05) between the normal RPE/choroid and retina -- *UCHL1, PFKP*, and *LPCAT1* (down-regulated in RPE/choroid vs. retina) and *PDPN, GAS1*, and *CST3* (up-regulated in RPE/choroid vs. retina) -- by real-time qPCR reactions run in triplicate on a subset of samples that were used for the RNA-Seq experiments. We confirmed the direction of effect for five of the six genes examined (data not shown). We were unable to detect *PFKP* expression in the RPE/choroid tissue and therefore this gene could not be validated. Additionally, we were able to replicate all of our top 20 genes from the normal RPE/choroid vs. normal retina by the Human Eye Integration data (https://eyeintegration.nei.nih.gov/). This database is a collection of healthy human RNA-Seq datasets generated from various studies of human eye tissue. To the best of our knowledge, no public database is yet available that contains gene expression data for the macular RPE/choroid tissues for different AMD clinical stages compared with normal macular RPE/choroid.

## Discussion

This is the first study using RNA-Seq to demonstrate genes, including a few of those previously associated with AMD risk, and lncRNAs are expressed differently in the macular retina compared to the macular RPE/choroid layers of eyes obtained within six hours of death from persons with intermediate or macular neovascularization or age-matched controls using a standardized protocol. Evaluation of known coding regions in previously reported GWAS loci demonstrated that significant ASE for *C3*, rs2230199, and *CFH*, rs1061170, occurred in the macula RPE/choroid for normal, and intermediate AMD while ASE for *CFH*, rs1061147, occurred in the macula RPE/choroid for normal, and intermediate and neovascular AMD. The protective variant for *CFB*, rs641153, only demonstrated ASE in the normal macular RPE/choroid. This is also underscored that these genes which function in the complement pathway were shown to be expressed statistically significantly higher in the normal macula RPE/choroid compared to the normal macula retina. Our comprehensive analysis demonstrated differential patterns of gene expression and splicing between diseased and non-diseased macula retina and RPE/choroid. Although the main pathology of AMD is affecting the macula area mainly, there is not many studies that compared the pathological changes occurring at the macula area (RPE/Choroidal) versus the inner retina, the relationship between the tissue types at given stages of disease is not entirely clear, leaving a big gap of knowledge related to this area [58–59]. It is clear that both the retina and RPE/choroid are important to AMD pathophysiology, it has been hypothesized that RPE cell function is more significantly related to AMD pathophysiology than retinal cell function [58, 60–62]. It is interesting to note that while we did not find the majority of the IAMDGC GWAS loci [5] differentially expressed between intermediate AMD and/or neovascular AMD, we did find that 75% of the genes were statistically significantly higher in the macular RPE/choroid compared to the macular retina thus underscoring the importance of tissue and geographic location in elucidating AMD etiology [62].

Of the known previously associated AMD loci [1, 10, 57, 63], none of them showed differential expression between disease states within the macular neural retina tissues. Of the 34 GWAS loci [5], expression differences between disease states were observed for *SPEF2* (between AREDS3 vs. Neo RPE) and *VTN* (between Neo vs. normal RPE/choroid). This suggests that variants in the gene regions showing heritable differences in AMD risk may influence the expression of tissue. A recent proteomic study on serums from patients with and without all types of late AMD (both neovascular and GA), found that *ZBPB, KREMEN2,* and *LINGO1*A were upregulated in diseased patients [64]. We found the expression of these genes to be significantly upregulated when comparing neovascular AMD or intermediate AMD with the normal macula RPE/choroid.

Absence of differential expression of the two genes previously reported to have the largest effect on AMD risk, *CFH* and *ARMS2/HTRA1*, is in agreement with studies demonstrating that *HTRA1* may influence AMD risk at the tRNA level [65–66]. Moreover, while *CFH* did not demonstrate differential gene expression between disease states in our study, *CFH* did demonstrate allele specific expression with both intermediate AMD and neovascular AMD depending on the SNP being interrogated. It may be this unequal expression of alleles at a given variant within the *CFH* gene to the disease pathophysiology of AMD. The mechanisms that underlie ASE are under active investigation and include epigenetics [67] In general, the differentially expressed genes reported here belong to pathways that have been shown to have important roles in AMD pathogenesis, including cholesterol and lipid metabolism (*DPP6, PON1, TIMD4*), mitochondrial regulation (*MTRNR2L1, TIMD4, IFI27, DISC1FP1*), and pro-inflammatory pro-macrophagic signaling in intermediate AMD [68–71]. Thus, our results also substantiate the possible contribution of macrophages in the late neovascular stage of the disease, as well as the contribution of mitochondrial dysfunction in the onset and progression of neurodegenerative diseases [68, 72]. Furthermore, *TCN2* regulates Vitamin B12 levels by encoding a protein that is part of the Vitamin B pathway, and was found in this study to be upregulated in those with intermediate AMD compared to normal macular RPE/choroid. Circulating low vitamin B12, high total homocysteine, and low folic acid levels are associated with AMD risk, suggesting that a defect in *TCN2* may reduce its ability to bind its partner proteins correctly, thus resulting in overexpression in the RPE/choroid and potentially a role in the subsequent development of large pathogenic drusen [73–78].

A trend was observed when comparing the top ten differentially expressed genes between each RPE comparison. In both the ‘intermediate AMD versus normal RPE/choroid’ and ‘neovascular AMD versus normal RPE/choroid’ comparisons, upregulation of *MTRNR2L1* was observed. Between both the ‘neovascular AMD versus normal RPE/choroid’ and ‘intermediate versus neovascular AMD RPE/choroid’ comparisons, three differentially expressed genes were shared, all expressed in opposite magnitudes (ie. upregulated in neovascular AMD and intermediate AMD, respectively; downregulated in normal and neovascular AMD, respectively): *RHO, RBP3,* and *RP51092A11.2.* Further investigation of these loci could prove useful for identifying novel biomarkers to differentiate between, as well as to track disease progression from, the intermediate form of AMD to the neovascular form.

We also observed differential expression of two distinct lncRNAs (*lncRNA AC00124.1* and *lncRNA RP11-240M16.1*) in the RPE/choroid of both intermediate and late-stage AMD patients in comparison to neovascular and normal eyes respectively, although in opposite directions (downregulated and upregulated, respectively). Since one lncRNA (*lncRNA AC000124.1*) was downregulated in intermediate AMD compared to neovascular AMD patients, it may be used as a biomarker for further validating disease progression from intermediate AMD to neovascular AMD. The lncRNA (*lincRNA RP11-240M16.*1) that we observed to be overexpressed in neovascular RPE/choroid compared to normal RPE/choroid may be a biomarker for late-stage disease. Further investigations could include comparing *lncRNA RP11-240M16.1* expression in neovascular versus GA AMD samples. If found to be differentially expressed between neovascular and GA forms of AMD, it be used as a biomarker to further distinguish between the two late-stage forms and could potentially contribute to pathogenesis. MicroRNAs may play an important role in AMD and represent a potential therapeutic target for treatment [79, 80]. The miRNA-146a was downregulated in RPE/choroid donor tissues from neovascular AMD subjects compared to controls. Previously, miRNA-146a was shown to be upregulated in the serum of patients with neovascular AMD [81–83]. Targets of miRNA-146a have been implicated in modulation of the immune response in endothelial tissue including negative regulation of complement factor H [84–86]. While the extent of involvement of the non-coding genome is still being investigates, miRANs, snoRNAs, and lncRNAs among non-coding RNA types are being found to have key roles in cellular homeostasis, and with disruption leading to human diseases such as cancer [87]. Our study is the first to evaluate these changes in the affected diseased tissue and suggest that there may be tissue differences in the manner in which miRNAs are expressed reinforcing the importance of examining expression in tissues affected directly by AMD. However further studies will need to be conducted in order to characterize fully the role of lncRNAs and miRNAs as biomarkers and determine their potential as therapeutic targets.

Many of our findings substantiate current ideas regarding AMD pathophysiology. Specifically, significant differential expression between disease states was observed for genes involved in lipid metabolism (*PON1*) [88] and inflammation (*TIMD4*, *PON1, DPP6*, *GPR123*)[55,89–90]. In addition, several of these genes have been associated with disease states hypothesized to co-occur more with AMD such as Alzheimer disease (AD) (*PON, SLC1A2, CPE, NME3*)[89–96] and epilepsy (*CHRNB2, SEZ6L2*) [97–99]. This suggests such diseases may have co-occurring pathophysiology as well as occurrence with AMD [99–101]. Our data also suggest a possible overlap in pathophysiology within the vitamin B12 synthesis pathway [101–102] and amyotrophic lateral sclerosis (ALS)[103–104]. Finally, we describe differential expression between intermediate and normal donor eyes of *CDH7,* a gene that was previously found to interact with smoking and be associated with AMD risk [52]. One gene, *MT2L1* (humanin), overlapped between RPE/choroid and retina, and was overexpressed in the intermediate stage of AMD compared to normal. Further studies on *MT2L1* have found it to serve a protective role in AD [105–107]. A similar circumstance has been noted for the inverse pattern of association of the APOE alleles: the ε4 allele increases risk of AD and the ε2 allele is protective, whereas the effects of these alleles on AMD risk are opposite [5,108–111].

Several genes and pathways were differentially expressed in comparisons of neovascular AMD with both intermediate AMD and normal macula, suggesting that clinical AMD phenotypes may represent a spectrum of molecular pathophysiology with a similar “core” of dysfunction with superimposed changes that dictate end-stage pathology. The majority of genes that were significantly differentially expressed in comparisons of normal macular tissue and intermediate AMD with end-stage neovascular disease participate in injury response (*SLC1A2, SCAMP5, RHO, PIWIL1, TM4SF2, SLC24A1, PTPRN, CPE, CHRNB2, CALB1*, and *CADM2*) and apoptosis (*CPE* and *PTPRN*) [110–116]. Genes identified in the phototransduction pathway (*ARR3*, *CNGA1*, *CNGA3*, *CNGB1*, *GNAT1*, *GNAT2*, *GNB1*, *GNB3*, *GNB5*, *GNGT1*, *GNGT2*, *GRK1*, *GUCA1A*, *GUCA1B*, *GUCA1C*, *GUCY2D*, *GUCY2F*, *OPN1LW*, *OPN1MW*, *OPN1SW*, *PDC*, *PDE6A*, *PDE6B*, *PDE6C*, *PDE6G*, *PDE6H*, *RCVRN*, *RGS9*, *RHO*, *SAG*) in macula of RPE/choroid of neovascular eyes, are important as retinal and choroidal neovascularization are the main causes of major visual impairment. Targeting molecules involved in photoreceptor apoptosis, has been suggested as an avenue for potential therapies to improve vision in neovascular AMD [118]. Pathway analysis of the overlapping DEGs from intermediate AMD vs normal AMD in the RPE/choroid and neovascular AMD vs normal AMD in the RPE/choroid showed the involvement of previously implicated GWAS hits TGF β signaling as well as BMP4 [119]. Thus, our findings may provide insight into the pathways modifying disease severity and or progression to neovascular end-stage disease.

Notch Signaling, WNT Beta Catenin Signaling, and TGF β signaling genes, as well as genes down-regulated in response to UV light, were down-regulated in the intermediate clinical stage macula compared to normal RPE/choroid macula. Of note, these pathways were not identified in the pathway analysis of differentially expressed genes in the retina comparing eyes from normal donors and persons with intermediate AMD (**Table 4**), and have all been implicated in AMD [57, 66, 120–133]. IL2/STAT5 signaling, IL6/JAK/STAT3 signaling, and the inflammatory response and epithelial mesenchymal transition were uniquely differentially expressed and upregulated in macular intermediate AMD compared to normal macular retina. Stat3 activation has previously been implicated in AMD pathophysiology, specifically in the neovascular subtype [134–139]. Taken together, these findings suggest that particular genes that function in the pathways of the RPE/choroid are protective in the development of the advanced forms of AMD while expression of STAT3 and/or other genes in inflammatory pathways in the retina may precede the development of late-stage AMD.

Within the RPE/choroid tissues, we identified several genes that were differentially expressed between normal donor eyes and both Intermediate AMD and neovascular AMD, but also, more importantly, gene expression profiles and pathways that were unique to these AMD stages. These observations are consistent with the idea that mechanisms governing the development of intermediate AMD are different from those influencing progression to neovascular AMD, [57, 140–146] and thus unique therapeutic interventions may be necessary to treat effectively these different forms of AMD. The histological changes that have been associated with neovascular AMD, including the new pathological growth of immature choroidal blood vessels under the retinal pigment epithelium (RPE) and/or in the subretinal space, moving towards the inner retina, transform or destroy the RPE layer resulting in a different histological arrangement of the retina (mixed RPE with retinal tissue) [147–149]. Therefore, understanding the different factors and genetic background involved in neovascularization and the resulting retinal changes are essential in the development of novel treatments for visual impairment. Our findings also underscore the importance of studying both macular tissue types to gain a full understanding of mechanisms leading to AMD. Future studies should focus on single cell and/or single nuclei from well characterized fresh tissue particularly.

In summary, these RNA-Seq and ASE experiments identified novel and established factors contributing to development of intermediate and neovascular AMD. Our results provide insights into underlying biological mechanisms that may differentiate the disease subtypes and into the tissues affected by the disease. It also expands upon previous RNA studies that demonstrated gene expression differences in affected tissues. Our results may provide insight into why some but not all individuals with intermediate AMD progress to the severe forms. If our results are confirmed in larger independent samples, differential gene expression could be implemented as an adjunct to a prognostic scheme (eg., blood). In order to make this a clinically useful tool, future studies will need to demonstrate equivalency in the same donors of pertinent gene expression patterns in eye tissue and other tissue that is easily accessible in living persons (e.g., blood).

## Supporting Information

**S1 Table. Significant differential gene expression.** Tables indicating all significant differential gene expression using a medium significance threshold (fold change > |1.5| and p < .05). There are separate tabs for each tissue and disease comparison.

